# A mix-and-match reverse genetics system for evaluating genetic determinants of orthobunyavirus neurological disease

**DOI:** 10.1101/2024.09.25.614932

**Authors:** Heini M. Miettinen, Matthew J. Abbott, Alyssa B. Evans

## Abstract

The encephalitic orthobunyaviruses have tri-segmented, negative sense RNA genomes and can cause severe neurological disease in humans, including La Crosse virus (LACV), which is the leading cause of pediatric arboviral encephalitis in the United States. However, little is known about the genetic factors that drive neuropathogenesis. Reverse genetics (RG) systems are valuable tools for studying viral genetics and pathogenesis. Plasmid-based cDNA RG systems are available for LACV, however the plasmid backbones are medium-copy number and have a propensity for recombination. We therefore generated a plasmid-based cDNA RG system for LACV utilizing a more stable and high-copy number plasmid backbone. Additionally, we created the first full RG systems for two closely related orthobunyaviruses, Jamestown Canyon virus (JCV), and Inkoo virus (INKV), which have differing reported disease incidences in humans and differing neuropathogenic phenotypes in mice compared to LACV. We compared wild type (wt) viruses with RG-derived viruses in human neuronal cells and in mice, and found that RG-derived viruses maintained the replication and neuropathogenic phenotypes of their wt counterpart. Additionally, we demonstrated that reverse genetics plasmids from different parental viruses can be readily mix-and-matched to generate reassortant viruses. This system provides a valuable genetic tool utilizing viruses with differing neuropathogenic phenotypes to investigate the genetic determinants of orthobunyavirus neuropathogenesis.

## INTRODUCTION

The Genus *Orthobunyavirus* contains many members capable of causing severe neurological disease in humans. These include closely related La Crosse virus (LACV), Jamestown Canyon virus (JCV), and Inkoo virus (INKV) in the California Serogroup (CSG; (1)). LACV is the leading cause of pediatric arboviral encephalitis in the USA and causes ∼100 neuroinvasive cases per year. The mortality rate of LACV is ∼1% and long-term sequelae including seizure disorders and cognitive defects are common in survivors. JCV is found in the USA and Canada and causes ∼50 neuroinvasive cases per year, primarily in adults. INKV is primarily found in Scandinavia and has been reported to cause a handful of neuroinvasive cases (2). Our previous studies on the neuropathogenesis of these viruses identified key differences in their ability to gain access to the brain from the periphery (neuroinvasion) and their ability to replicate and cause damage and disease in the brain (neurovirulence) (3, 4). We found that LACV was neuroinvasive with high neurovirulence, JCV was not neuroinvasive but with high neurovirulence, and interestingly, INKV was neuroinvasive but with low neurovirulence. The viral determinants that mediate neuropathogenesis are not well understood.

Orthobunyaviruses have three negative-sense, single-stranded RNA genome segments: L, M, and S. The L segment encodes the RNA-dependent RNA polymerase (L protein; (5). The M segment encodes a polyprotein with the two envelope glycoproteins (Gn and Gc) and a nonstructural protein (NSm) of uncertain function (6, 7). The S segment encodes the nucleocapsid protein (N) which forms protective ribonucleoprotein complexes with the RNA genome, and nonstructural protein S (NSs), which is encoded within the N reading frame and results from leaky ribosomal scanning (7, 8). NSs is an interferon (IFN) antagonist (9) and the only known virulence factor for LACV.

Previous studies of orthobunyavirus genetic determinants of neuropathogenesis have largely relied on generating reassortant viruses using highly passaged attenuated viruses with unknown mutations (10, 11). More recent studies have utilized LACV reverse genetics (RG) systems to investigate the effect of targeted mutations, gene deletions, or ORF replacements in LACV neuropathogenesis (12, 13, 14). While this approach provides critical insights into viral genetics, a limitation of this strategy to investigate viral determinants of neuropathogenesis is that mutations may disrupt protein functions in multiple ways. Viral proteins often serve multiple roles during infection, which can confound results and make teasing out mechanism from viral gene mutations and knockouts difficult.

To develop an RG system that keeps viral proteins unchanged, we leveraged the key differences in neuropathogenic phenotypes between LACV, JCV, and INKV to create full cDNA plasmid-based RG systems for each virus. We showed that RG-derived viruses maintained their replication and neuropathogenic phenotypes in human neuronal cells and mice compared to their wild type (wt) counterpart. We also showed we can generate replication competent RG-derived reassortant viruses by mix-and-matching L, M, and S RG plasmids between the different viruses. With this system we generated a higher fidelity LACV RG system, as well as the first reported full RG systems for JCV and INKV. This system provides a valuable genetic tool utilizing wild type versions of viral genes from viruses with disparate neuropathogenic phenotypes to investigate the viral determinants of orthobunyavirus neuropathogenesis.

## METHODS

### Cells and viruses

Gibco reagents were used unless otherwise specified. BHK-21/BSR/T7/5 (BSR/T7) cells were maintained in GMEM supplemented 2mM L-Glutamine solution, 1x MEM Amino Acid Solution, and 10% FBS (Atlas Biologicals EF-0500-A), and every other passage with 1mg/ml Geneticin. Vero cells (ATCC) were maintained in DMEM supplemented with 10% FBS and 1x penicillin-streptomycin. SH-SY5Y cells (ATCC) were maintained in 1:1 ratio of EMEM (ATCC) and F-12K (ATCC) supplemented with 10% FBS and 1x pen-strep. LACV (human 1978), JCV (61V2235), and INKV (SW AR 83-161) were gifted by Dr. Karin Peterson (NIAID). Intermediate stocks of wild type (wt), RG, and RG-reassortant viruses were generated in Vero cells with media supplemented with 25µg/ml of Plasmocin (Invivogen NC9886956) and harvested at 80% CPE at 2-4 dpi, then final stock viruses made in normal media.

### Generation of reverse genetics plasmids and viruses

Plasmids were constructed using our wt virus stocks and sequences of LACV human 1978, JCV 61V2235, and INKV SW-AR-83-161. All plasmids were generated in the pMK vector backbone. Plasmids containing the full coding regions and UTRs of JCV and INKV M and S segments flanked by a T7 promoter and hepatitis delta virus ribozyme as previously described (12, 13) were synthesized by Invitrogen GeneArt in the pMK vector backbone to generate reverse genetics plasmids pJCV-M, pJCV-S, pINKV-M, and pINKV-S. pMK plasmid vectors encoding the LACV L UTRs and portions of the INKV 3’ and 5’ coding regions or the JCV L UTRs and portions of the JCV 3’ and 5’ coding regions flanked by the T7 promoter and ribozyme were synthesized by GeneArt (pINKV-L_linker and pJCV-L_linker, respectively).

To make the LACV-RG plasmids, viral RNA from LACV-wt was isolated via Qiagen’s Viral RNA Mini Kit and cDNA generated using the iScript Select cDNA synthesis kit (Bio-Rad) and virus-specific primers (Eurofins; supplemental materials). Full-length cDNAs of LACV L, M, and S segments were made using Q5 High-Fidelity 2x Master Mix (New England Biolabs) using segment-specific primers with complementary 5’ overhangs with the vector backbone. The vector backbone was amplified using T7 promoter and ribozyme-specific primers. DNA assembly of the fragments was done using NEBuilder HiFi DNA Assembly Master Mix (New England Biolabs) at 50°C for 15 min. Two µl of the assembly mix was transformed into homemade chemically competent Stbl3 *E. coli* to generate pLACV-L, pLACV-M, and pLACV-S.

pINKV-L was generated using INKV-wt isolated RNA. cDNA was made using SuperScript IV VILO master mix (Invitrogen), and the coding region of INKV L amplified by PCR and purified using the Wizard SV Gel and PCR Clean-up System (Promega). The INKV L fragment and the pINKV-L_linker backbone were digested with *Aat* II and *Bsr* GI (New England Biolabs), purified, ligated with T4 DNA ligase (New England Biolabs) and transformed into homemade chemically competent Stbl3 cells. To generate pJCV-L, four fragments with 21-25 nucleotide overlaps encoding the JCV L segment were synthesized (GenScript). pJCV-L_linker was amplified using 2x Q5 high-fidelity master mix, digested with *Dp*n I to remove the template DNA and gel purified. DNA assembly was carried out using the NEBuilder HiFi DNA Assembly Master Mix for 60 min at 50°C. 2 µl assembly mix was transformed into 50 µl New England Biolabs 5-α chemically competent *E. coli* according to the manufacturer’s instructions. All plasmids were sequenced-verified by Plasmidsaurus and sequences analyzed with MacVector 18.5.1.

Transfections with the RG plasmids were done by plating BSR/T7 cells at 1×10^5^ cells per well in a 6-well plate ∼24 hrs prior to transfection. Transfections were performed using TransIT-LT1 (Mirus MIR2304) according to manufacturer’s instructions with the following modifications: 0.5 µg of an L, M, and S plasmid and 9 µl TransIT-LT1 reagent was used per transfection mix. Supernatants were harvested and clarified at 48-72 hpt to recover RG viruses.

### Replication kinetics

Replication kinetics assays were performed as previously described (3). Briefly, SH-SY5Y cells were plated at 1×10^5^ cells per well in 96-well plates. The next day, cells were inoculated in triplicate at MOI=0.01, incubated 1 hour at 37C, washed twice, and media replenished. Supernatants were harvested and clarified at 1-96 hpi, then titered on Vero cells. Three independent experiments were performed.

### Virus titers

Vero cells were plated in 24-well plates at ∼1.3×10^5^ cells per well. The next day, virus was serially diluted 10-fold (replication kinetics) or 5-fold (virus stocks) and plated in duplicate in 200µl per well. Plates were incubated 1 hour at 37C, then overlayed with 1.5% carboxymethyl cellulose (ThermoSci A18105.36) in MEM (Gibco 12360-038). Plates were fixed with 10% formaldehyde and stained with 0.35% crystal violet at 4 dpi.

### Mice

All mouse experiments were approved by Montana State University’s Institutional Animal Care and Use Committee under protocol number 2023-238-IA and were performed in accordance with NIH guidelines. Neuroinvasive disease was evaluated by inoculating weanling mice (21-23 days of age) intraperitoneally with 1×10^5^ PFU virus. Neurovirulence was evaluated by inoculating adult mice (>6 weeks of age) intranasally with 1×10^4^ PFU virus. All mice were followed for signs of neurological disease including ataxia, seizures, paralysis, and circling, or other endpoint criteria including moribundity, and were immediately humanely euthanized. Any surviving mice were humanely euthanized by 25 dpi.

### Statistical analyses

All statistical analyses were performed in GraphPad Prism v. 10.2. Replication kinetics were evaluated by two-way ANOVA with Sidak’s multiple comparisons test to determine differences between wt and RG viruses at any time point. Additional simple linear regression of slopes was performed for the log phase of viral growth at 6 to 72 hpi to compare 1) wt vs RG versions of the same virus and 2) all viruses compared to LACV-wt reference. Survival curves were analyzed via Mantel-Cox and Gehan-Breslow-Wilcoxon tests. For all analyses, p-values ≤0.05 were reported as significant.

## RESULTS & DISCUSSION

### Generation of reverse genetics-derived viruses

We utilized a strategy that has been used for other LACV reverse genetics (RG) systems with individual DNA plasmids for each of the L, M, and S segments under the control of a T7 promoter with a hepatitis delta virus ribozyme at the 3’ end (12, 13). Previous LACV RG systems have utilized the pBR322 vector backbone. However, this medium-copy number plasmid has a propensity for recombination which can frequently result in plasmids with full or partial duplications, making targeted genetic changes a challenge. We therefore used a more stable high-copy number plasmid, pMK, as the backbone vector for our RG plasmids. The full viral segments from LACV, JCV, and INKV L, M, and S segments were generated using our wild-type (wt) stock viruses (virus-wt) reference sequences. We used a variety of cloning and DNA synthesis techniques to generate plasmids pLACV-L, pLACV-M, pLACV-S, pJCV-L, pJCV-M, pJCV-S, pINKV-L, pINKV-M, and pINKV-S. All plasmids encode the full coding sequence and UTRs from each individual viral segment, with the exception of pINKV-L. This plasmid contains the full L coding sequence from INKV with the LACV L segment UTRs, due to insufficient sequence information on the INKV L segment UTRs. Plasmids were sequenced and compared to wt reference sequences. pLACV-L had a single nonsynonymous mutation, pJCV-L had two nonsynonymous and one synonymous mutation, pINKV-L had two nonsynonymous mutations, and pINKV-M had two nonsynonymous and one synonymous mutation (Table 1).

**Table 1.** Sequence comparison of plasmids with reference.

| **Table 1.** Sequence comparison of plasmids with reference |  |  |  |  |  |  |
| --- | --- | --- | --- | --- | --- | --- |
| Segment | nucleotide |  |  | amino acid |  |  |
|  | pos | LACV-wt | LACV-RG | pos | LACV-wt | LACV-RG |
| L | 2357 | C | T | 766 | L | L |
|  | pos | JCV-wt | JCV-RG | pos | JCV-wt | JCV-RG |
| L | 4975 | G | T | 1641 | D | Y |
|  | 4978 | C | T | 1642 | P | S |
|  | 5523 | C | T | 1823 | F | F |
|  | pos | INKV-wt | INKV-RG | pos | INKV-wt | INKV-RG |
| L | 2486 | G | A | 829 | R | Q |
|  | 2808 | G | A | 936 | M | I |
| M | 2340 | G | A | 763 | R | K |
|  | 2794 | G | A | 914 | Q | Q |
|  | 3545 | C | T | 1165 | L | L |
| Nonsynonymous |  |  |  |  |  |  |

Infectious virus was generated via transfection of the L, M, and S plasmids for each virus in BHK-21 cells that constitutively express T7 polymerase. Supernatants containing virus were collected and titered on Vero cells. To compare viral titers between wt and RG viruses, viruses were inoculated at MOI=0.01 on Vero cells and supernatants were harvested at ≥80% CPE. Both the wt and RG versions of each virus grew to similar titers (Figure 1), suggesting that RG-derived viruses maintain the replication and infectivity phenotypes of their wt virus counterpart.

**Figure 1.**
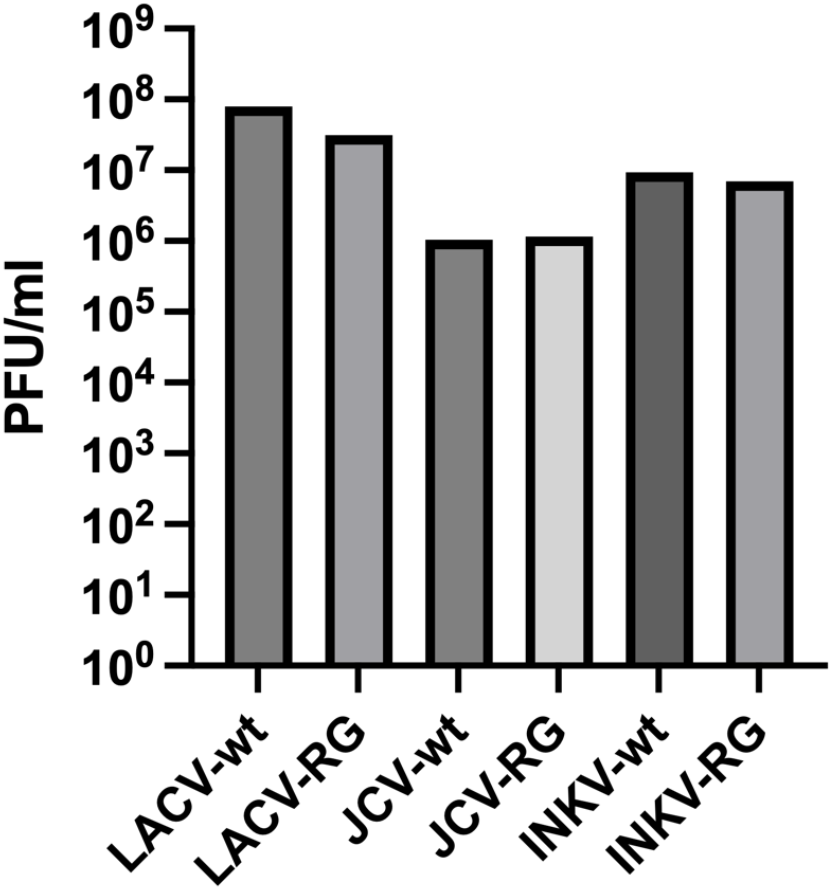
Comparison of wt and RG viral titers. Vero cells were infected with wt and RG-derived viruses at MOI=0.01. Viruses harvested at ≥80% CPE, then titered on Vero cells.

### RG viruses maintain wild type replication phenotypes in neuronal cells *in vitro*

To further evaluate whether RG-derived viruses replicated similarly to their wt counterpart, we performed replication kinetics assays. We have previously used SH-SY5Y neuroblastoma cells to compare the replication kinetics of wt LACV, JCV, and INKV and found that JCV and INKV replicated slower than LACV (3). We therefore evaluated the replication kinetics of our wt and RG-derived viruses in SH-SY5Y cells. We inoculated cells with each virus at an MOI=0.01 and collected supernatants at 1-96 hpi. Supernatants were titered on Vero cells.

Overall, the replication kinetics of wt vs RG viruses of the same virus were similar, although there were some minor time point titer differences, none of these were significantly different via two-way ANOVA (Figure 2; Table 2). All wt and RG-derived viruses of the same virus reached similar peak titers on the same day (Figure 2, Table 2). Linear regression analysis found no significant differences in viral replication between wt or RG-derived viruses of the same virus (Table 2). However, similar to our previous comparisons of wt LACV, JCV, and INKV, there were differences between virus strains in replication kinetics (Table 2). Both JCV and INKV viruses had significantly slower replication kinetics compared to LACV, consistent with our previous studies (3). Together, these results indicate that the RG-derived viruses maintain their wt replication phenotypes *in vitro*.

**Table 2.** Analysis of Replication Kinetics.

| Table 2 Analysis of Replication Kinetics |  |  |  |  |  |
| --- | --- | --- | --- | --- | --- |
|  | ANOVA p-values§ | LR p-value† | LACV-wt† | (PFU/ml) | (hpi) |
| LACV-wt | 0.63-0.98 | 0.282 | ref | 7.98 x 10<sup>7</sup> | 48 |
| LACV-RG |  |  | 0.282 | 1.23 x 10<sup>8</sup> | 48 |
| JCV-wt | 0.14-0.99 | 0.779 | **0.032** | 5.98 x 10<sup>7</sup> | 96 |
| JCV-RG |  |  | **0.013** | 4.13 x 10<sup>7</sup> | 96 |
| INKV-wt | 0.74-0.99 | 0.877 | **0.007** | 4.08 x 10<sup>7</sup> | 96 |
| INKV-RG |  |  | **0.007** | 3.28 x 10<sup>7</sup> | 96 |
| §2-way ANOVA with Sidak's multiple comparisons test. |  |  |  |  |  |
| †Linear regression analysis of slopes covering the log phase of viral growth, 6-72 hpi. |  |  |  |  |  |
| Statistically significant |  |  |  |  |  |

**Figure 2:**
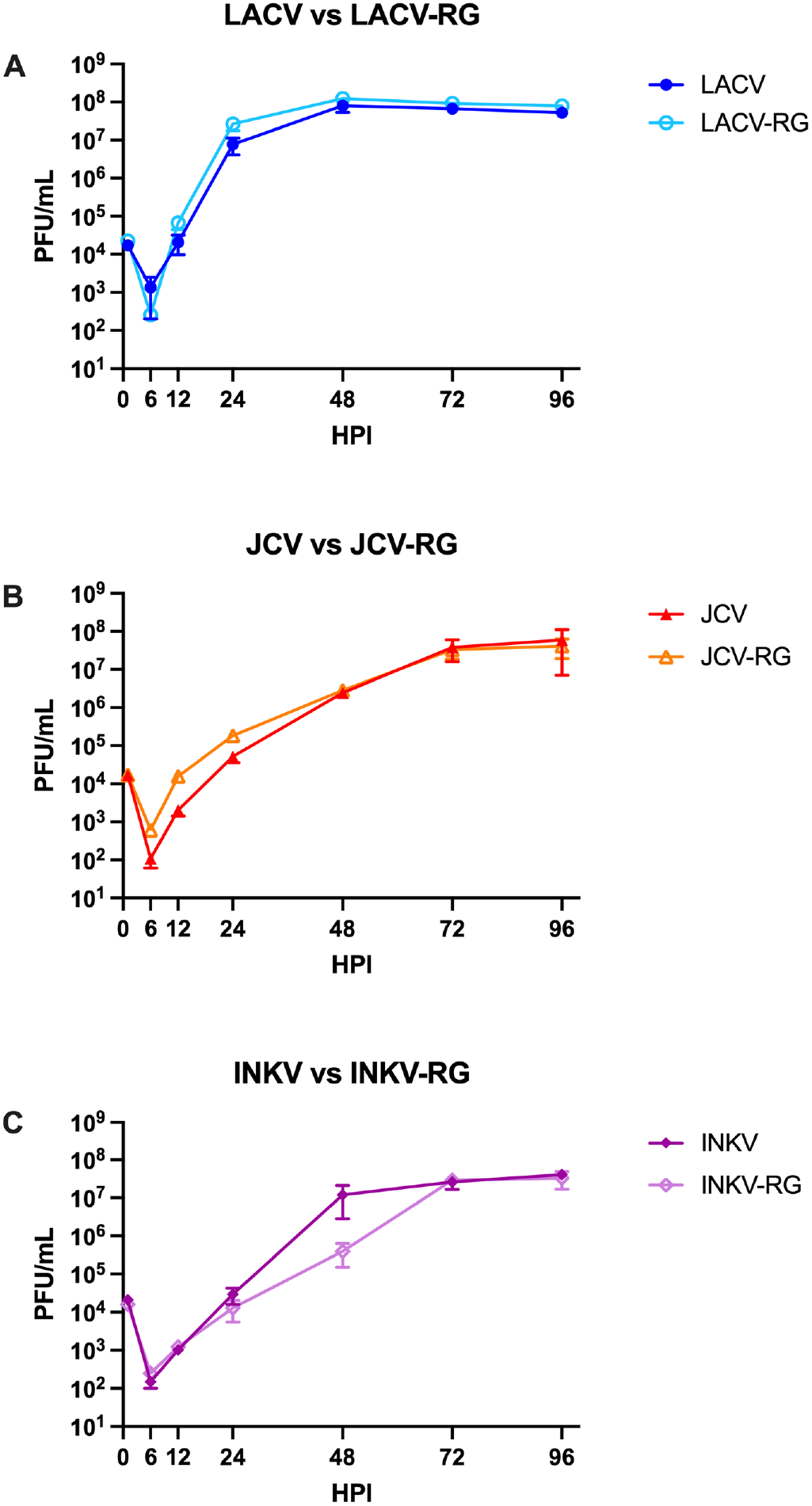
Replication kinetics of wt vs reverse genetics viruses in SH-SY5Y cells. Data represents three independent experiments titered on Vero cells.

### RG viruses maintain wild type neuropathogenic phenotypes in mice *in vivo*

We next evaluated if the RG-derived viruses maintained their neuropathogenic phenotypes compared to their wt counterpart in mice. In our previous comparisons of LACV, JCV, and INKV in their ability to induce neuroinvasive disease after peripheral inoculation in weanling mice, we found that LACV caused disease in 100% of mice whereas JCV and INKV did not cause neuroinvasive disease in any mice (3). We therefore inoculated weanling mice IP with 1×10^5^ PFU of the viruses to assess the development of neuroinvasive disease. Neurological signs included ataxia, paralysis, circling, and seizures. Both LACV-wt and LACV-RG induced neuroinvasive disease in 100% of mice at 4-6 dpi (Figure 3A). JCV-wt and JCV-RG did not cause neuroinvasive disease in any of the mice. INKV-RG did not cause neuroinvasive disease in any mice, however INKV-wt caused neuroinvasive disease in one out of twelve mice (Figure 3A). While we had not previously observed a mouse develop neuroinvasive disease from INKV, we have previously shown that INKV enters the brains of weanling mice in the absence of disease (3, 4). Therefore, it may not be surprising that occasionally an INKV-inoculated mouse may develop neuroinvasive disease. However, there was no significant difference between any wt and RG virus of the same strain via simple survival curve analysis.

**Figure 3.**
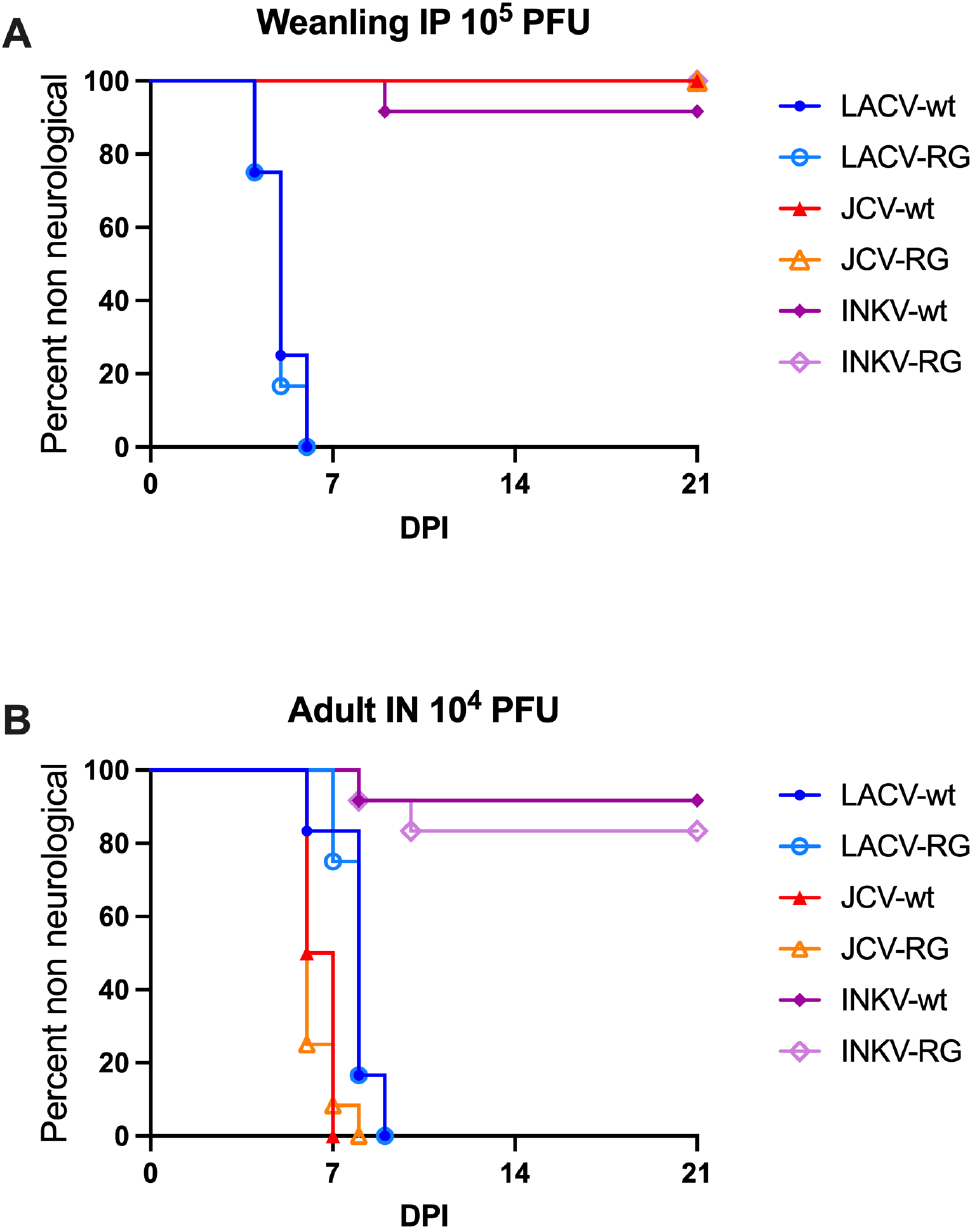
Comparison of wt and RG virus pathogenicity in mice. A) Weanling mice were inoculated intraperitoneally with 1×10^5^ PFU virus. B) Adult mice were inoculated intranasally with 1×10^4^ PFU virus. Mice were followed for neurological signs of disease. n=11-12

To evaluate neurovirulence phenotypes, we inoculated adult mice intranasally with 1×10^4^ PFU and followed them for neurological signs. At this route and dose, we have previously shown that LACV and JCV induced disease in 100% of mice, whereas INKV induced disease in ∼15-20% of mice (3). Consistent with previous results, LACV-wt caused neurological disease in 100% of mice at 6-9 dpi, and LACV-RG caused disease in 100% of mice at 7-9 dpi (Figure 3B). JCV-wt caused disease in 100% of mice at 6-7 dpi and JCV-RG caused neurological disease in 100% of mice at 6-8 dpi (Figure 3B). INKV-wt and INKV-RG induced neurological disease in one and two mice out of twelve, respectively (Figure 3B). There was no significant difference between any wt and RG virus of the same strain via simple survival curve comparison analysis.

Together, these results demonstrate that RG-derived viruses maintain the neuropathogenic phenotypes of their wt counterpart, making RG-derived viruses an accurate tool for comparing differences in orthobunyavirus neuropathogenesis.

### RG-generated reassortant viruses are successfully recovered from mix-and-match plasmid transfections

As proof-of-principal that we can generate reassortant viruses using this reverse genetics system, we next performed mix-and-match transfections between LACV & INKV, LACV & JCV, and JCV & INKV to generate M segment reassortant viruses LILV-RG, LJLV-RG, IJIV-RG, and JIJV-RG, which are summarized in Figure 4A. We evaluated stock titers of the RG-derived reassortant viruses in Vero cells in the same manner as the wt and RG viruses (Figure 1). Consistent with the replication kinetics results, wt and RG-derived viruses of the same strain had very similar titers (Figure 4B). The RG-reassortant viruses were more variable in titer than the parental viruses, but still reached titers of ∼5×10^5^ – 2×10^7^ PFU/ml, depending on the virus (Figures 1, 4B). These results demonstrate that infectious virus can be recovered by mixing and matching L, M, and S plasmids from different parental viruses.

**Figure 4.**
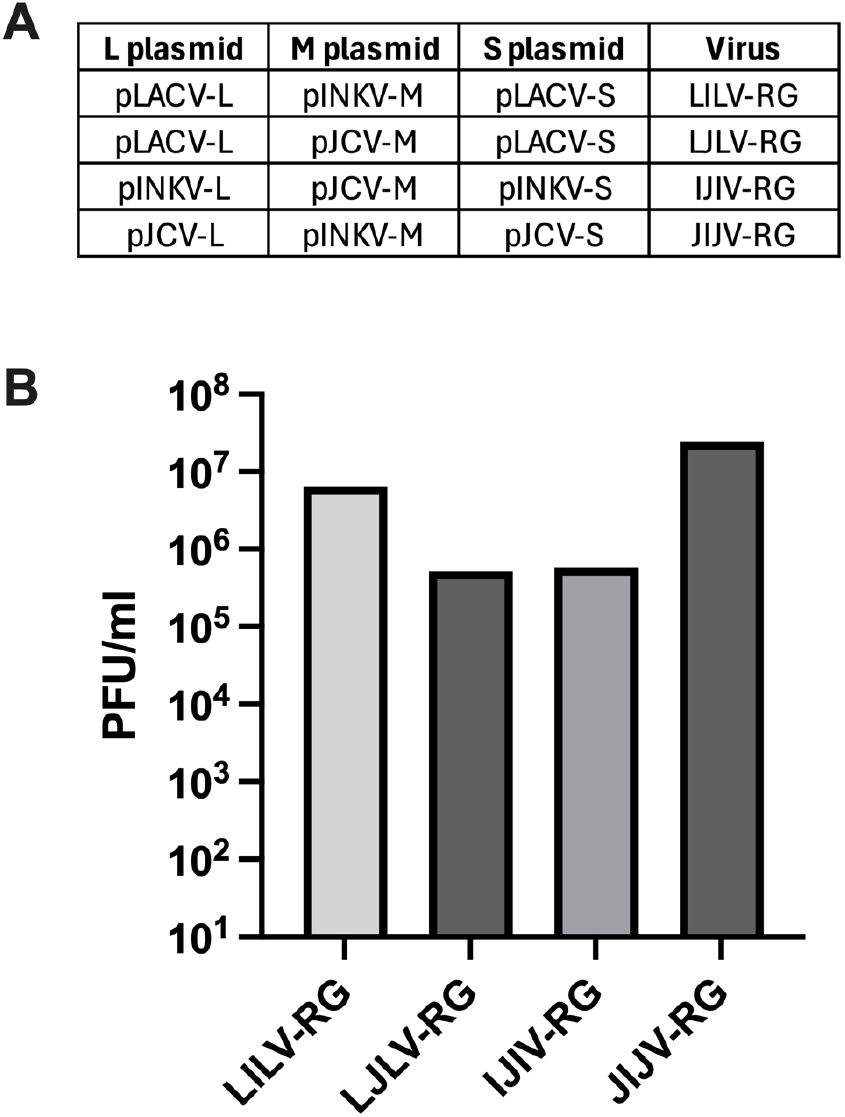
Generation of Mix-and-Match Reassortant-RG viruses. A) Table of the plasmids used for mix-and-match transfections in BSR/BHK/T7 cells to recover the reassortant viruses. B) Vero cells were infected with wt and RG parental viruses and RG-derived reassortant viruses at MOI=0.01. Viruses harvested at ≥80% CPE via visual inspection, then titered on Vero cells.

### Conclusions

Determining the underlying genetic factors that mediate orthobunyavirus neuropathogenesis is critical to understanding orthobunyavirus neurological disease and developing effective therapeutics and vaccines. To facilitate genetic comparisons between orthobunyaviruses with differing neuropathogenic phenotypes, we generated reverse genetics systems for LACV, JCV, and INKV. The three RG-derived viruses maintained the neuropathogenic phenotypes of their wt counterparts both *in vitro* and *in vivo*. Furthermore, plasmids can be readily mix-and-matched between parental viruses to generate reassortant viruses, making these an ideal tool for genetic studies of orthobunyavirus molecular pathogenesis.

## ACKNOWLEDGEMENTS

We thank Dr. Karin Peterson for providing critical reagents for this project. We thank the animal technicians at the MSU Animal Resource Center who helped facilitate the mouse work.

